# A paradoxical relationship between mitochondrial calcium regulation and retinal ganglion cell degeneration after axon damage

**DOI:** 10.64898/2026.05.13.724793

**Authors:** Sean McCracken, Minglei Zhao, Kyler Squirrell, Christopher Zhao, Saman Behboodi Tanourlouee, Michelle Aum, Philip R. Williams

**Affiliations:** John F. Hardesty, MD Department of Ophthalmology and Visual Sciences, Washington University School of Medicine, St. Louis, MO 63110, USA; Department of Neuroscience, Washington University School of Medicine, St. Louis, MO 63110, USA; Hope Center for Neurological Disorders, Washington University School of Medicine, St. Louis, MO 63110, USA

## Abstract

Retinal ganglion cells (RGCs) degenerate in optic neuropathies like glaucoma and traumatic optic nerve injury leading to irreversible vision loss. Higher levels of homeostatic Ca^2+^ and canonical Ca^2+^ regulated signaling promote RGC survival in animal models of glaucoma and optic nerve injury. Mitochondrial dysfunction is also a hallmark of degenerating neurons, including RGCs. Here, we investigate the intersection of mitochondrial function, Ca^2+^ homeostasis, and cellular resilience by performing an optic nerve crush model of RGC degeneration while monitoring and manipulating mitochondrial Ca^2+^ levels (mito-Ca^2+^). We find that mito-Ca^2+^ is predicative of RGC survival in that surviving RGCs are enriched for higher homeostatic mito-Ca^2+^ levels. Mitochondrial dysfunction was observed where mito-Ca^2+^ was reduced in RGCs after injury, regardless of survival. We then examined the importance of higher mito-Ca^2+^ in surviving RGCs by altering mito-Ca^2+^ levels and Ca^2+^ transit using pharmacological and AAV-mediated approaches. Paradoxically, treatment to decrease mito-Ca^2+^ increased survival to ONC. We then manipulated mito-Ca^2+^ permeability by altering the expression levels of the mitochondrial calcium uniporter (MCU) pore forming subunit that allows Ca^2+^ to enter mitochondria from the cytoplasm. Overexpressing MCU reduced RGC survival to injury, while shRNA knockdown of MCU increased RGC survival. These results reveal a complex relationship between mito-Ca^2+^ and RGC degeneration and suggest that well-surviving RGCs may be under chronic mitochondrial stress due to higher homeostatic mito-Ca^2+^ levels.

## INTRODUCTION

Retinal ganglion cells (RGCs) are the final output neurons of the retina that possess long-distance projecting axons susceptible to insult and injury. Axon damage *via* mechanical injury of the optic nerve leads to retrograde cell death and irreversible vision loss in optic neuropathies including glaucoma and traumatic optic nerve injury (Carelli et al., 2004; Quigley, 2016). Preserving and restoring vision in these optic neuropathies requires RGC neuroprotection, and there are currently no treatments for vision maintenance and repair that target RGCs directly. Therefore, understanding mechanisms that underly RGC death is critical for preservation and restoration of vision.

Dysregulation of intracellular Ca^2+^ occurs in a variety of neurodegenerative conditions (Anwar, 2016; Bezprozvanny, 2010; Gandhi et al., 2009; Tu et al., 2006) and likely contributes to RGC degeneration in optic neuropathies (Crish and Calkins, 2011; Li et al., 2025). Under normal physiological conditions, Ca^2+^ homeostasis is tightly regulated to ensure proper cellular function. However, in retinal degeneration, pathological stressors disrupt this Ca^2+^ balance, which can trigger a cascade of detrimental events, including mitochondrial dysfunction (Zhang et al., 2023), activation of Ca^2+^-dependent proteases like calpains (Kobayashi-Otsugu et al., 2020; Vu et al., 2022), and initiation of apoptosis (Nickells, 1999). In contrast to a detrimental role for Ca^2+^ overload driving neurodegeneration, we recently identified a diversity in homeostatic cytoplasmic Ca^2+^ set-points in RGCs that correlate with survival following optic nerve crush (ONC) (McCracken et al., 2023). Intriguingly, we found that RGCs that maintained higher homeostatic Ca^2+^ levels in the cytosol were more likely to survive ONC, and that lowering Ca^2+^ levels reduced the survival of high baseline Ca^2+^ RGCs (McCracken et al., 2023). Since intracellular Ca^2+^ is regulated over multiple organellar sub-compartments and can directly influence the function of these organelles, we were motivated to determine the relationship between RGC degeneration and Ca^2+^ homeostasis in specific Ca^2+^ regulating organelles.

Intracellular Ca^2+^ regulation is a dynamic process that spans the cytosol, mitochondria and endoplasmic reticulum (ER). The ER is a critical Ca^2+^ store that amplifies Ca^2+^ signaling through Ca^2+^ induced Ca^2+^ release. Dysfunction of RGC ER-Ca^2+^ dynamics occurs in mouse models of glaucoma leading to prolonged electrophysiological responses to light (Shiga et al., 2024). This ER-Ca^2+^ dysfunction can be corrected by Serca2 overexpression, which protects RGCs (Shiga et al., 2024). Mitochondrial Ca^2+^ (mito-Ca^2+^) is important for many neuronal functions including cytosolic Ca^2+^ buffering, intracellular signaling, recovery after stimulation, regulation of ATP production, and stress responses (Amrapali Vishwanath et al., 2026; Matuz-Mares et al., 2022; Walters and Usachev, 2023). Mito-Ca^2+^ overload, in particular, is a key factor in neuronal apoptotic death as it impairs mitochondrial energy production and exacerbates oxidative stress (Giorgi et al., 2012; Kowalczyk et al., 2021). However, the precise mechanisms that regulate cellular Ca^2+^ homeostasis, and how they might impact survival during RGC neurodegeneration have not been investigated.

In this study, we used *in vivo* two-photon microscopy to investigate homeostatic organellar Ca^2+^ levels in mouse RGCs and their dynamics in response to ONC. We did not find any relationship between RGC survival and homeostatic or dynamic ER-Ca^2+^ levels. However, similar to cytosolic Ca^2+^ (McCracken et al., 2023), RGCs with higher homeostatic mito-Ca^2+^ levels were more likely to survive during degeneration. In contrast to cytosolic Ca^2+^, altering mito-Ca^2+^ homeostasis through genetic or pharmacologic approaches targeting MCU showed a clear inverse relationship between mito-Ca^2+^ and RGC survival, where reducing mito-Ca^2+^ improved RGC survival and increasing mito-Ca^2+^ transit decreased survival. These findings show that although well-surviving RGCs maintain higher homeostatic mito-Ca^2+^ levels, this trait is actively detrimental to their survival after a degenerative stimulus.

## RESULTS

We investigated Ca^2+^ levels across different intracellular domains using a fluorescence resonance energy transfer (FRET)-based Ca^2+^ biosensor, Twitch2b, as previously described (McCracken et al., 2023; Thestrup et al., 2014). Twitch2b was expressed in RGCs using an intersectional viral and transgenic approach consisting of VGlut2-Cre transgenic mice and adeno-associated virus serotype 2 (AAV2) packaged with a Cre-dependent Twitch2b expression cassette. Cytoplasmic Ca^2+^ (cyto-Ca^2+^) was measured using Twitch2b without a localization signal (T2b), while an N-terminal cytochrome c oxidase 8 fusion was used to target Twitch2b to the inner mitochondrial matrix (mito-T2b; (Witte et al., 2019)). To compare cyto- and mito-Ca^2+^ levels, AAV2-FLEX-T2b and AAV2-FLEX-mito-T2b were injected into 3-4 week old VGlut2-Cre mice and RGCs were then imaged 2-3 weeks later with *in vivo* trans-pupillary 2-photon microscopy (Wang et al., 2021). In line with previous measurements that found varied cyto-Ca^2+^ levels across RGCs (McCracken et al., 2023), we observed diverse homeostatic mito-Ca^2+^ levels (Figure 1A). Mito-Ca^2+^ levels were, on average, similar to cyto-Ca^2+^ levels while displaying a slightly less variable distribution. (Cyto-T2b YFP/CFP = 1.20 ± 0.43, mito-T2b = 1.21 ± 0.30 (mean ± SD unless otherwise indicated), p = 0.95 MWU test; Figure 1B). Mito-Ca^2+^ levels were relatively consistent within individual RGCs imaged 4-6 days apart (R = 0.45; Figure 1C-E).

**Figure 1.**
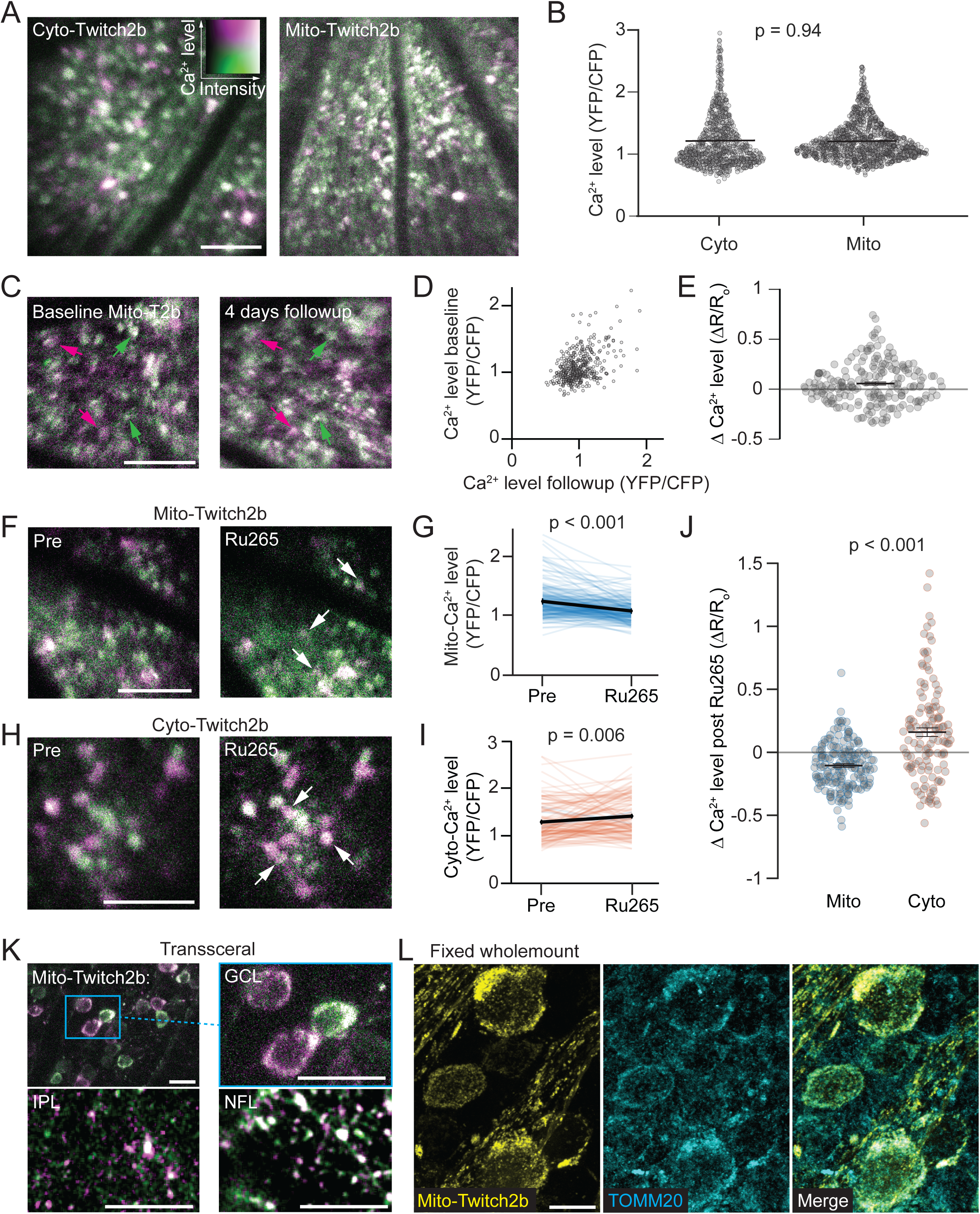
*In vivo* RGC Mitochondrial Ca^2+^ levels. (A) Example *in vivo* 2-photon max intensity projections (mip) of cytoplasmic (Cyto, left) and mitochondrial (Mito, right) Twitch-2b expressed in retinal ganglion cells (RGCs) of VGlut2-Cre mice. (B) Swarm plots of homeostatic cyto- and mito-Ca^2+^ levels, as measured by cpVenus (YFP) to mCerulean (CFP) FRET ratios, MWU test (cyto-T2b n = 868 RGCs from 14 retinas and 8 mice, mito-T2b n = 1066 RGCs from 19 retinas and 15 mice). (C) Average intensity projections (aip) of mito-T2b *in vivo* of the same RGCs at baseline and 4 days later. Magenta arrows indicate RGCs with higher baseline mito-Ca^2+^ and green arrows indicate RGCs with lower mito-Ca^2+^ levels. (D) Scatterplot of mito-Ca^2+^ levels pre and 4 day follow-up timepoints. R = 0.45, (n = 205 RGCs from 5 retinas and 3 mice). (E) Swarm plots of change in mito-Ca^2+^ levels from baseline to 4 day follow-up replotted from panel (D). (F) Example aips of mito-T2b at baseline and 10 min following intravitreal Ru265 injection. White arrows indicate RGCs with reduced mito-Ca^2+^. (G) Line graphs of mito-Ca^2+^ levels before and 10 min after Ru265 injection. Individual RGCs are in blue, and mean is in black (n = 175 RGCs from 4 retinas and 4 mice). (H) Example aips of cyto-T2b at baseline and 10 min after intravitreal Ru265. White arrows indicate RGCs with increased cyto-Ca^2+^. (I) Line graphs of cyto-Ca^2+^ levels before and 10 min after Ru265 injection. Individual RGCs are in orange, and mean is in black (n = 123 RGCs from 4 retinas and 4 mice). (J) Swarm plot comparing changes in Ca^2+^ levels after Ru265 between cyto- and mito-T2b. Black bars are mean +/- SEM replotted from panels (G and I). (K) Example aips of KCNG4-Cre mito-T2b acquired with *in vivo* transscleral imaging. GCL = ganglion cell layer, IPL = inner plexiform layer, NFL = nerve fiber layer. Box indicates magnified region. (L) Representative confocal mips of fixed retinal wholemounts showing endogenous mito-T2b (yellow) and TOMM20 (cyan) immunostaining. Scale bars = 100 μm.

### Validation of mito-T2b

To validate that mito-T2b specifically measures mito-Ca^2+^, we assessed an array of compounds that block mitochondrial Ca^2+^ import by injecting them intravitreally and recording mito-T2b responses (data not shown). We found that an analog to the Ruthenium Red family of compounds, Ru2(μ-N)(NH3)8Cl2]Cl3, trans, trans-Octaamminedichloro-μ-nitridodi-Ruthenium(3+) trichloride, Nitride, ruthenium complex (Ru265), was most effective at decreasing mito-Ca^2+^ specifically *in vivo*. Injection of Ru265 (1 μL, 2 mM in 50% dimethylsulfoxide (DMSO) and 50% PBS) caused a strong reduction in RGC somatic mito-Ca^2+^ (Pre 1.24 ± 0.02 vs. Post = 1.08 ± 0.14, p < 0.001 MWU test; Figure 1F-G) that was sustained for 86 +/- 12 min (mean +/- SEM; data not shown) and was dose dependent (Supplemental Figure 1). To confirm this response was specific to mitochondria, we then tested the effect of Ru265 injection on cyto-Ca^2+^. Intravitreal injections of 2 mM Ru265 in mice expressing cyto-T2b demonstrated increased Ca^2+^ levels (Pre = 1.28 ± 0.04 vs. Post = 1.41 ± 0.04, p = 0.006 MWU test; Figure 1H-I), suggesting that Ru265 specifically reduces mito-Ca^2+^ levels. The increase in cyto-Ca^2+^ levels after Ru265 injection was in strong contrast with the reduction in mito-Ca^2+^ levels (mito-Twitch2b ΔR/R_o_ = -0.11 ± 0.01; cyto-Twitch2b ΔR/R_o_ = 0.16 ± 0.03; p < 0.001, MWU test; Figure 1J). The unexpected elevations in cyto-Ca^2+^ levels after Ru265 injection are likely due to the drug being dissolved in DMSO since vehicle alone elevated both cyto- and mito-Ca^2+^, especially at earlier observation points (Supplemental Figure 2). These data indicate that Ru265 delivery specifically elevates mito-Ca^2+^ despite the DMSO vehicle causing Ca^2+^ elevations.

To confirm mito-T2b localization to the mitochondria, we imaged low density RGCs at high resolution *in vivo* using transscleral 2-photon microscopy (Alarcon-Martinez et al., 2020) in albino (B6(Cg)-Tyr c-2J /J)) KCNG4-Cre mice injected with AAV-FLEX-mito-T2b, which restricts mito-T2b to αRGCs (Duan et al., 2015). 3-D image volumes were obtained from the nerve fiber layer (NFL) through the ganglion cell layer (GCL) and inner plexiform layer (IPL) to capture mitochondria from all regions of RGCs (Figure 1K). Distribution of mito-T2b appeared as expected for mitochondrial localized signal, with an absence of diffuse contiguous labeling in RGC arbors. A dense network of mito-T2b was present in the RGC somata but excluded from nuclei. Mito-T2b labeled structures in the NFL were non-contiguous and mostly elongated, whereas in the IPL they were shorter and more pill-shaped to spheroid in appearance. We also used high resolution confocal scanning of fixed wholemount retinas with RGCs expressing mito-T2b in VGlut2-Cre mice to validate mitochondrial specificity. We observed similar distributions of mito-T2b as *in vivo* with dense, nuclear excluded networks in the soma and non-contiguous structures in axons and dendrites. Importantly, mito-T2b signal was colocalized with immunostaining for Translocase of Outer Mitochondrial Membrane 20 (TOMM20), a canonical mitochondrial marker ((Buso et al., 2019; Pitt and Buchanan, 2021) Figure 1L).

Some evidence of mis-localization of the mito-T2b biosensor was observed in RGCs that expressed mito-T2b strongly and for an extended time period (Supplemental Figure 3). In wholemount preparations of mito-T2b labeled retinas that were counterstained with TOMM20, most RGCs had mito-T2b that overlapped with TOMM20 signal, but a small fraction showed partial mis-localization with signal present diffusely in the cytosol. Fewer RGCs showed complete mis-localization where T2b filled the cell aside from a few distributed vacuoles. Importantly, the rate of mis-localized mito-T2b was correlated with expression levels, and RGCs with mis-localized mito-T2b accumulated over time after AAV-FLEX-mtio-T2b delivery (<30 dpi: 95 ± 0.08 % localized, >60 dpi: 86 ± 2.7% localized; Supplemental Figure 3E-I). Thus, to be sure the vast majority of our RGCs represented true mito-Ca^2+^ levels, we restricted our experiments to samples that were from 2-4 weeks after AAV-FLEX-mito-T2b injection.

### High baseline mitochondrial Ca^2+^ levels are associated with well surviving RGCs

We previously reported a relationship with high cyto-Ca^2+^ levels and RGC survival after ONC axon injury (McCracken et al., 2024). To investigate the role of mito-Ca^2+^ in this phenotype, first we examined if mito-Ca^2+^ levels were higher in well-surviving RGC subtypes. We imaged mito-Ca^2+^ levels *in vivo*, then performed post hoc immunostaining on the imaged regions (Figure 2A) using antibodies to label αRGCs (SPP1, (Duan et al., 2015)), and intrinsically photosensitive RGCs (ipRGCs) (TBR2, (Chen et al., 2021)), which are both generally well-surviving RGC subtypes (Tran et al., 2019). The intersection of these two antibodies labels ON-sustained αRGCs/M4 RGCs, which is one of the most well-surviving RGC types (Schmidt et al., 2014; Tran et al., 2019). SPP1+ RGCs demonstrated higher mito-Ca^2+^ levels than SPP1- RGCs (SPP1 + vs. SPP1 -, 1.36 ± 0.05 vs. 1.16 ± 0.02, p < 0.001, MWU test), and TBR2+ RGCs displayed higher mito-Ca^2+^ levels than TBR2- RGCs (TBR2 + vs. TBR2 -, 1.35 ± 0.05 vs. 1.18 ± 0.02, p < 0.001, MWU test). Furthermore, the ON-sustained αRGCs (SPP1+/TBR2+) had the highest mito-Ca^2+^ levels of all groups examined (Figure 2B). Since the SPP1+/TBR2+ double positive RGCs demonstrated such high mito-Ca^2+^ levels, removing them from the TBR2+ group led to a loss of significance for TBR2+ cells when comparing these RGC types to the general population that were non-stained(TBR2+/SPP1- vs. All stains-, p = 0.10; Figure 2B). Therefore, mito-Ca^2+^ levels in ON-sustained α/M4 RGCs possess mito-Ca^2+^ levels higher than most RGCs, including some of their related ipRGC types. Lastly, we attempted to confirm that αRGCs as a cohort had higher mito-Ca^2+^ levels than the general RGC population by measuring mito-T2b in KCNG4-Cre as compared to VGlut2-Cre transgenic mice. We did not confirm our immunostaining results as mito-Ca^2+^ levels were not different between VGlut2-Cre or KCNG4-Cre RGC populations (Figure 2C-D). Taken together, the well-surviving ON-sustained αRGC demonstrates significantly higher mito-Ca^2+^ levels than other RGCs.

**Figure 2.**
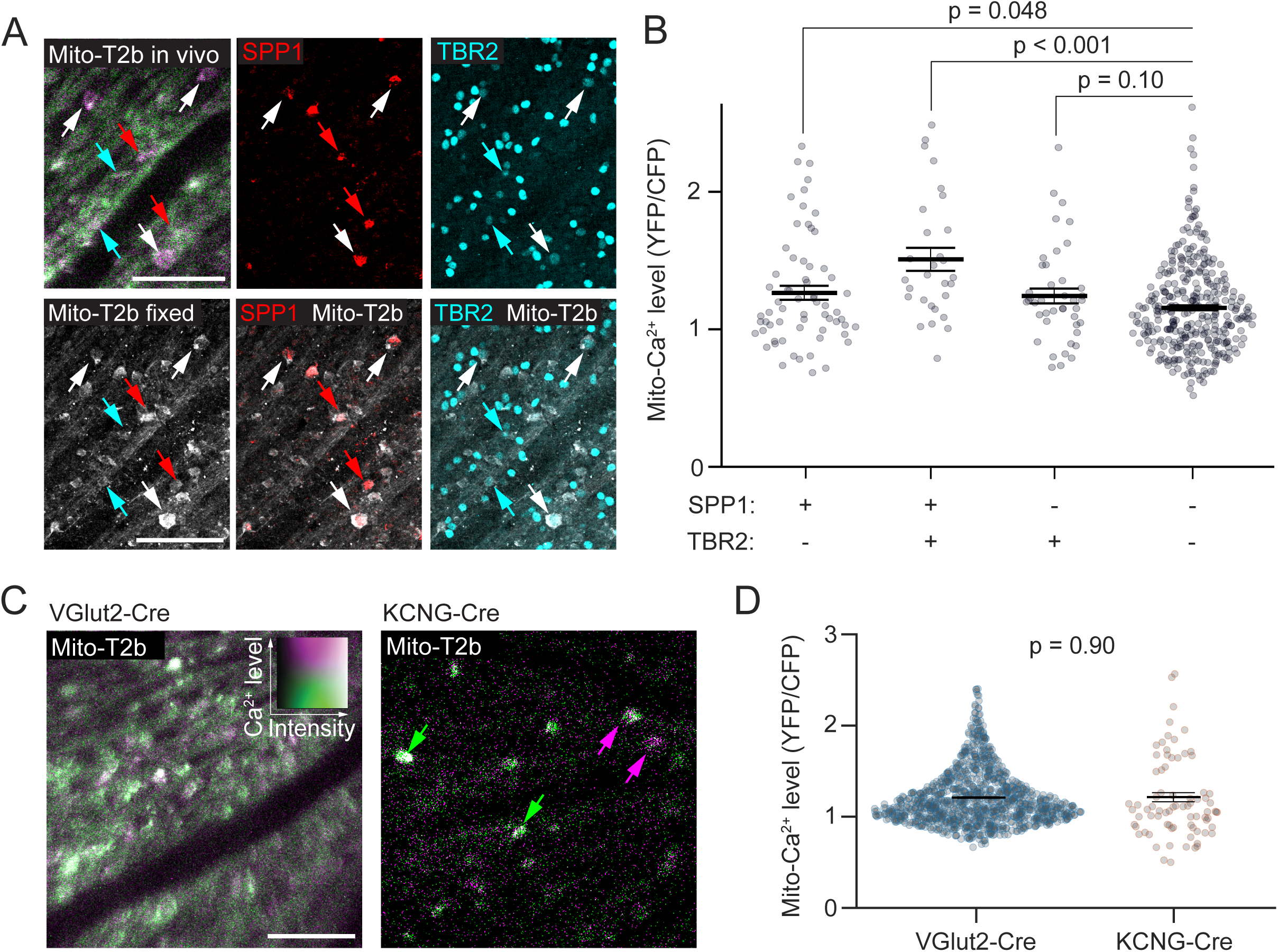
Mitochondrial Ca^2+^ levels are higher in some well-surviving RGC subtypes. (A) Example *in vivo* aips of mito-T2b from VGlut2-Cre transgenic mouse and the same imaged region in fixed retinal wholemount immunostained for SPP1 (red, αRGCs) and TBR2 (cyan, ip-RGCs). Mito-T2b is greyscale. Arrows represent cells identified as positive for the different stains: cyan = TBR2 only, red = SPP1 only, and white = SPP1 / TBR2 positive. (B) Swarm plots of mito-Ca^2+^ levels in individual RGCs positive for indicated RGC markers. Means are labeled with a black bar +/- SEM (n = 479 RGCs from 6 retinas and 6 mice). (C) Representative aips of mito-T2b expression in VGlut2-Cre (all RGCs) and KCNG4-Cre transgenic mice (αRGCs). (D) Swarm plots of mito-Ca^2+^ levels from VGlut2-Cre and KCNG4-Cre transgenic mice (n = 79 RGCs from 6 retinas and 3 mice). Scale bars = 100 μm.

To directly examine how mito-Ca^2+^ levels are related to RGC survival, we tracked RGCs that expressed mito-T2b with longitudinal *in vivo* imaging after ONC. We first obtained baseline images of mito-Ca^2+^, performed ONC 2 to 4 days later, and then obtained subsequent *in vivo* images every 2 days from 4 to 14 days post ONC (dpc) when the majority of RGC death occurs (Figure 3A-B; (Li et al., 2020)). We observed that surviving RGCs had higher baseline mito-Ca^2+^ levels than dying RGCs (surviving 1.37 ± 0.04 vs. dying: 1.21 ± 0.01, p < 0.001, MWU test; Figure 3C). To determine the time course of RGC degeneration as related to differences in mito-Ca^2+^ levels, we grouped RGCs into “high” and “low” baseline mito-Ca^2+^ using the upper and lower halves of mito-T2b ratios, determined within each imaging field. We observed a consistent trend for high-Ca^2+^ RGCs to survive better than lower Ca^2+^ cells, with a significant difference at 4, 8, and 10 dpc (Figure 3D). The higher rate of RGC survival suggests that higher homeostatic mito-Ca^2+^ levels are beneficial for RGCs after axon injury.

**Figure 3.**
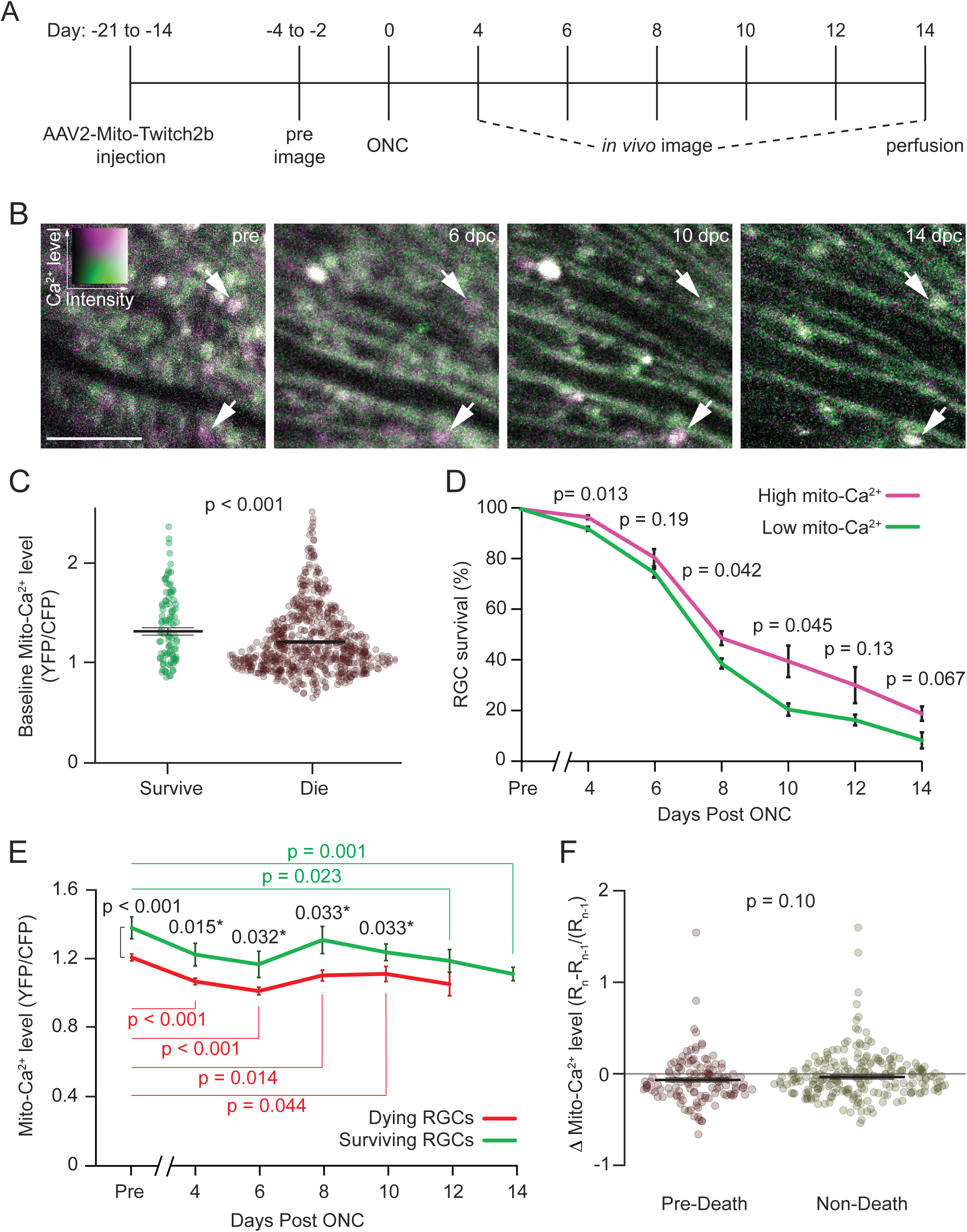
Surviving RGCs have higher homeostatic mitochondrial Ca^2+^ levels. (A) Timeline depicting *in vivo* imaging and ONC experiments. (B) Example *in vivo* 2-photon aips of mito-Ca^2+^ expressing RGCs before ONC and at the indicated days post ONC (dpc). (C) Swarm plots of pre-injury mito-Ca^2+^ levels in RGCs that survived (green) or died (red) following ONC. Means +/- SEM are labeled with black bars (survive n = 100 RGCs, die n = 669 RGCs from 5 retinas and 3 mice,). (D) Survival curves of RGCs with high (magenta) or low (green) pre-injury mito-Ca^2+^ levels. Error bars are SEM (n = 295 RGCs from 3 retinas and 3 mice). (E) Dynamic analysis of mito-Ca^2+^ levels in RGCs that survived (green) or died (red) during *in vivo* imaging time course. Statistics comparing surviving RGCs (green) or dying RGCs (red) to their pre-injury mito-Ca^2+^ levels, or surviving vs. dying RGCs (black) (from n = 3 retinas and 3 mice including 45 +/- 4.7 surviving RGCs and 119 +/- 38 dying RGCs per sample). (F) Dynamics of mito-Ca^2+^ levels between 2-day time points either prior to death (pre-death) (n = 126 RGCs from 3 retinas and 3 mice) or from all other timepoints (non-death) (n = 225 RGCs from 3 retinas and 3 mice). Scale bar = 100 μm.

### Mito-Ca^2+^ levels are reduced after ONC injury

Since influxes of Ca^2+^ into the mitochondrial compartment drive neuronal death in many neurodegenerative disorders (Calvo-Rodriguez et al., 2020; Jurcau and Jurcau, 2023; Muller et al., 2018; Supnet and Bezprozvanny, 2010; Verma et al., 2017), we next tracked mito-Ca^2+^ dynamics in surviving and dying RGCs after ONC. Instead of mito-Ca^2+^ elevations, we observed a decrease in mito-Ca^2+^ levels that were exacerbated over time after ONC (Figure 3E). Despite this change, surviving RGC mito-Ca^2+^ levels remained higher than those of dying RGCs at most timepoints (Figure 3E). To examine if overload of mitochondrial Ca^2+^ drives RGC death, as has been observed in other neurodegenerative conditions (Calvo-Rodriguez et al., 2020; Muller et al., 2018; Verma et al., 2017), we evaluated mito-Ca^2+^ changes that occurred over the observations immediately prior to death (pre-death) and compared these to any other time-point to time-point changes in mito-Ca^2+^ levels after which RGCs did not die (non-death). While we did not see any significant difference in pre-death mito-Ca^2+^ dynamics vs. non-death (Figure 3F), we did observe decreases in mito-Ca^2+^ over time in both surviving and dying associated intervals corresponding with the dynamic observations of an overall reduction in mito-Ca^2+^ chronically after ONC (Figure 3E). This contrasts with cytosolic Ca^2+^ levels, which are relatively stable after ONC (McCracken et al., 2023). Thus, mito-Ca^2+^ import from the cytosol is actively reduced in response to degeneration. Together, these data demonstrate that mito-Ca^2+^ is reduced after ONC in both surviving and dying RGCs over time, suggesting the mito-Ca^2+^ dysregulation occurs in injured RGCs. It is possible that brief elevations in mito-Ca^2+^ occur just prior to RGC death, but given the large number of dying RGCs we observed (670 cells), this is unlikely. Overall, mito-Ca^2+^ dysregulation manifests as a reduction in mito-Ca^2+^ levels, in contrast to most observations in other neuronal populations that would link mito-Ca^2+^ overload to degeneration.

### Endoplasmic Reticulum Ca^2+^ is not correlated with RGC survival

The ER is a major intracellular Ca^2+^ store and dysregulation of ER-Ca^2+^ occurs in many forms of neurodegeneration (Schrank et al., 2020; Shiga et al., 2024). To directly examine ER-Ca^2+^ in RGCs, we first performed baseline *in vivo* imaging of VGlut2-Cre transgenic mice injected with AAV2-FLEX-ER-T2b, which used an N-terminal fusion of the mouse Vh-chain ER-signal peptide for ER localization (ER-T2b; (Witte et al., 2019)). ER-Ca^2+^ levels were significantly higher than cytoplasmic or mito-Ca^2+^, in accordance with well-documented high Ca^2+^ levels in the ER (Alvarez and Montero, 2002; Lipscombe et al., 1988) (ER-T2b = 2.73 ± 0.77, cyto-T2b = 1.20 ± 0.43, mito-T2b = 1.21 ± 0.30, p < 0.001 compared to cyto-Ca^2+^ or mito-Ca^2+^, MWU test; Supplemental Figure 4A-B). To determine if there is a relationship between ER-Ca^2+^ and RGC survival after ONC, we tracked ER-Ca^2+^ in RGCs every 2 days for 2 weeks after ONC. Unlike cytoplasmic or mito-Ca^2+^, there was no difference in homeostatic ER-Ca^2+^ between dying or surviving RGCs (ER-T2b surviving = 2.54 ± 0.10 vs. dying = 2.50 ± 0.06, p = 0.55; MWU Test; Supplemental Figure 4C-D). Taken together, it does not appear that ER-Ca^2+^ homeostasis is a major driver of native survival to RGC degeneration.

### Intermittent reductions of mito-Ca^2+^ with Ru265 protects RGCs from ONC

Since we observed that higher homeostatic and dynamic mito-Ca^2+^ levels were associated with RGC survival, and well-surviving ON-sustained αRGCs demonstrated higher mito-Ca^2+^ levels, we wanted to test if higher mito-Ca^2+^ was causative of RGC survival. Therefore, we lowered mito-Ca^2+^ using Ru265, which we determined to lower mito-Ca^2+^ specifically for more than an hour at 2 mM dosages (Figure 1). We performed repeated intravitreal injections of Ru265 starting immediately after ONC and injecting every three days until termination of experiments at 14 days post ONC (Figure 4A). We tested Ru265 at three different concentrations (20 μM, 200 μM, and 2 mM) compared to vehicle control (50% DMSO in PBS) and examined RGC survival in these treatments by immunostaining wholemounts for the RGC marker RNA-binding protein with multiple splicing (RBPMS) (Rodriguez et al., 2014). Although higher mito-Ca^2+^ was observed in well-surviving RGCs, we observed that Ru265 injections increased RGC survival in a dose dependent manner (Figure 4B-C). Repeated injections of Ru265 or vehicle without ONC did not cause significant RGC death, while other control injections of PBS alone or sham injections did not show any differences compared to vehicle after ONC (Supplemental Figure 5). These data indicate a curious relationship between mito-Ca^2+^ and survival where surviving RGCs demonstrate higher homeostatic mito-Ca^2+^ levels (Figures 2-3), yet manipulating mito-Ca^2+^ to lower concentrations is protective after injury. This relationship contrasts with cytoplasmic Ca^2+^, where higher cytoplasmic Ca^2+^ is observed in well-surviving RGCs, and reducing cytoplasmic Ca^2+^ reduces RGC survival (McCracken et al., 2023).

**Figure 4.**
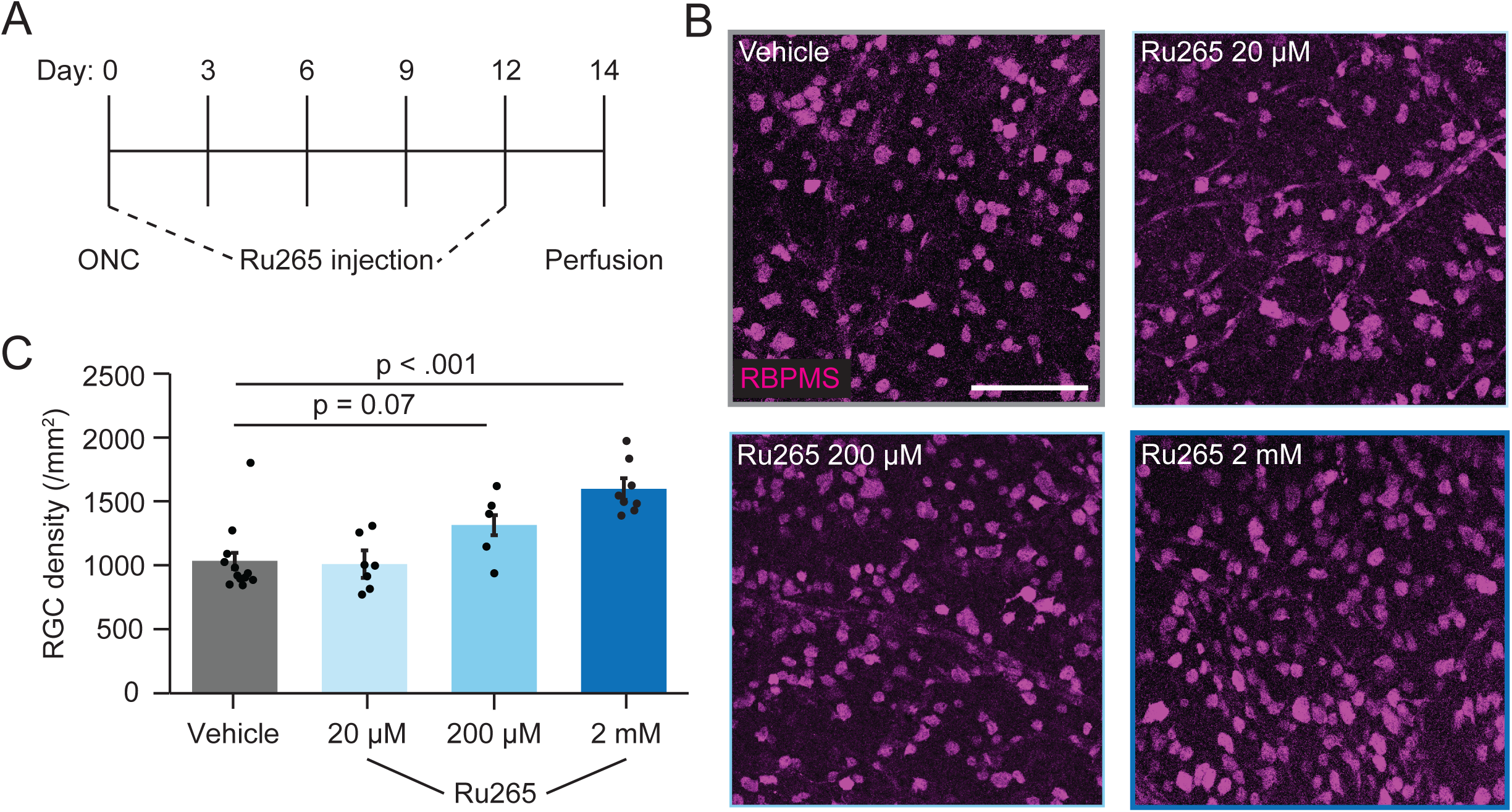
Longitudinal Ru265 injections protect RGCs from ONC. (A) Timeline depicting ONC and Ru265 injection experiments. (B) Representative mips of retinal wholemounts immunostained with RBPMS at 14 dpc with indicated treatments. (C) Quantification of RBPMS densities from indicated experimental conditions. Points are individual RGCs, bars are mean +/-SEM, MWU test (vehicle n = 12 retinas from 12 mice, 20 μM Ru265 n = 7 retinas from 7 mice, 200 μM Ru265 n = 5 retinas from 5 mice, 2 mM Ru265 n = 8 retinas from 8 mice). Scale bar = 100 μm.

### Targeting the mitochondrial calcium uniporter to promote RGC survival after ONC

Since Ru265 transiently alters mito-Ca^2+^, it may be possible that transient yet large fluctuations of mitochondrial permeability to Ca^2+^ alter mitochondrial function and influence RGC survival differently to chronic alterations of mito-Ca^2+^ permeability. Thus, we attempted to stably shift homeostatic mito-Ca^2+^ using a gene therapy approach targeting the mitochondrial calcium uniporter (MCU), an inner mitochondrial membrane protein that is required for Ca^2+^ import into the mitochondria (Baughman et al., 2011; De Stefani et al., 2011; Kirichok et al., 2004). To do this, we used AAV2 vectors to overexpress (MCU-OX) or knockdown MCU (shMCU (Qiu et al., 2013)), with AAV2-FLEX-MCU-T2A-mCherry and AAV2-U6-shMCU-CAG-mCherry viruses respectively, injected into VGlut2-Cre transgenic mice. We confirmed an increase in MCU expression in MCU-OX retinas by immunostaining for MCU. RGCs transfected with MCU-OX virus, as inferred by mCherry reporter signal, had stronger MCU immunofluorescence than neighboring RGCs without the presence of the mCherry reporter (Supplemental Figure 6). Native MCU expression was relatively low as assayed by immunofluorescence in accordance with transcriptional data (Tran et al., 2019). As such, we did not observe a decrease in MCU expression in shMCU treated samples given the low baseline expression, but there was a trend for reduced MCU expression in cells with the highest levels of mCherry reporter expression (Supplemental Figure 6).

To examine the effects of MCU manipulations on RGC survival, we injected mice with AAVs driving either MCU-OX or shMCU (Figure 5A) and compared against control AAV2-FLEX-mCherry virus injected eyes primarily in eye matched control samples to reduce variability. Three weeks later, ONC was performed and two weeks post-ONC, mice were perfused and RGC survival was measured by RBPMS immunostaining (Figure 5B). Consistent with Ru265 results, decreasing MCU expression with shMCU protected RGCs from degeneration (shMCU RBPMS+ density 1077 ± 143 vs. mCherry controls 608 ± 90, p = 0.02 MWU Test; Figure 5C-D,F). Conversely, MCU-OX decreased RGC survival after ONC (MCU-OX RBPMS+ density 547 ± 55 vs. mCherry controls 871 ± 84, p = 0.007, MWU Test; Figure 5E-G). These experiments support our findings using Ru265 in that reducing mito-Ca^2+^ protects RGCs from degeneration.

**Figure 5.**
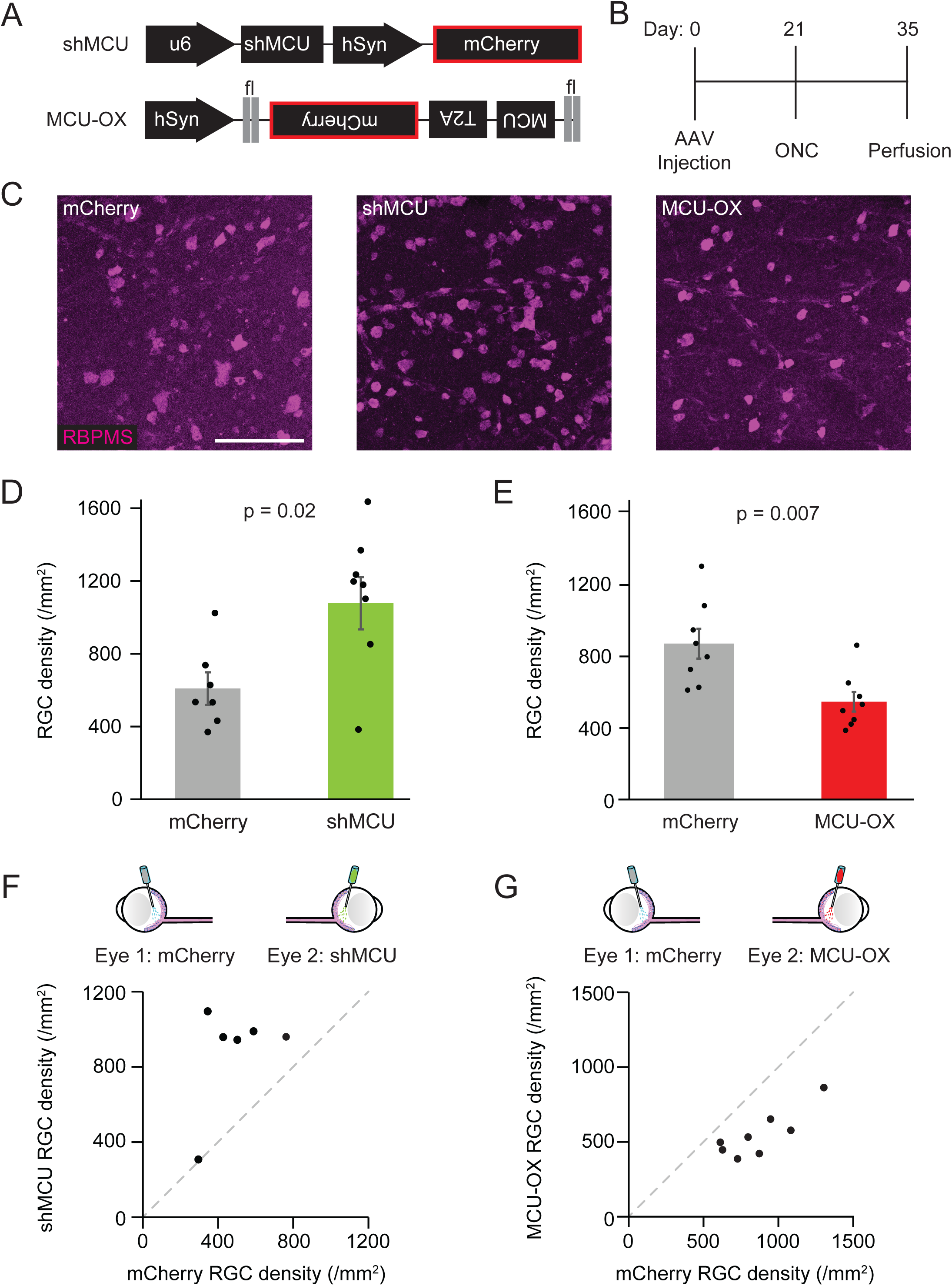
Mitochondrial Calcium Uniporter expression regulates RGC survival after ONC. (A) Schematic illustrating AAV constructs used to knockdown (shMCU) or overexpress (MCU-OX) mitochondrial calcium uniporter protein (MCU). (B) Time course of AAV injection and ONC experiments. (C) Representative mips of fixed retinal wholemounts immunostained for RBPMS at 14 dpc. (D-E) Quantification of RBPMS+ cell density in mice with intravitreal injections of indicated AAV MCU gene expression manipulations compared to littermate AAV-mCherry treated controls collected 14 dpc. Points indicated individual eyes bars indicate mean +/- SEM, MWU test (D: shMCU n = 8 retinas from 8 mice and mCherry n = 7 retinas from 7 mice, 6 eye matched; E: MCU-OX n = 8 retinas from 8 mice and mCherry n = 8 retinas from 8 mice, all eye matched). (F-G) Paired-eye comparison of RGC density following indicated AAV treatment or mCherry control. Samples are also included in panels D-E. Dashed line is y = x. Scale bar = 50 μm.

Since chronically manipulating mito-Ca^2+^ could itself lead to neurodegeneration (Singh et al., 2022), we examined if RGCs degenerated in response to MCU manipulation without ONC. Two weeks after sham ONC, neither MCU-OX nor shMCU demonstrated reductions in RGC density (Supplemental Figure 7), showing that over the time course of ONC experiments, baseline degeneration is not driven by MCU manipulations. However, by 8 weeks after sham ONC, MCU-OX and shMCU both induced significant RGC death (sham mCherry control RGC density 4655 ± 109 RGCs/mm^2^; MCU-OX 2762 ± 236 RGCs/mm^2^, p<.001; shMCU 4025 ± 181 RGCs/mm^2^, p = 0.025; MWU test vs mCherry; Supplemental Figure 7). Thus, while chronically increasing mito-Ca^2+^ transit with MCU overexpression was toxic in the long term, we do not expect this toxicity alone to underlie reduced RGC survival after ONC. Furthermore, although shMCU treatment was protective in ONC, a strong model of RGC degeneration, long term reduction in MCU expression appears detrimental to RGC health.

## DISCUSSION

Our study reports a paradoxical relationship between homeostatic mitochondrial Ca^2+^ levels and RGC survival in response to axon injury. Using *in vivo* imaging to measure Ca^2+^ levels in mitochondrial and ER compartments, we demonstrated that well-surviving RGCs are enriched for high homeostatic mito-Ca^2+^ levels, whereas ER-Ca^2+^ was not correlated with survival outcome. This observation is in accordance with our previous experiments that showed high cytoplasmic Ca^2+^ levels are a feature of well-surviving RGCs (McCracken et al., 2023). However, we found an inverse relationship between the higher mito-Ca^2+^ phenotype of well-surviving RGCs and the impact of altering mito-Ca^2+^ levels on RGC survival. Using pharmacological and gene therapy based manipulations, we show that lowering mito-Ca^2+^ is protective (Figures 4&5). Conversely, MCU overexpression leads to more RGC death (Figure 5). These results indicate that not all features of well-surviving RGCs drive this resilient phenotype, and some may in fact be detrimental to survival.

RGCs with higher circuit activity and cytoplasmic Ca^2+^ are well-surviving, and lowering circuit activity or intracellular Ca^2+^ reduces RGC survival, especially those with high homeostatic Ca^2+^ levels (McCracken et al., 2023; Tran et al., 2019; Zhang et al., 2019b). Similarly, treatments that elevate retinal circuit activity, and thus stimulate intracellular Ca^2+^ signaling, protect RGCs from ONC (Li et al., 2016; Lim et al., 2016; Zhang et al., 2019b). Although CaMKII signaling is strongly protective of RGCs in ONC and glaucoma models (Guo et al., 2021), in a previous study, we did not find any relationship between homeostatic cytoplasmic Ca^2+^ and CaMKII signaling in healthy or injured RGCs (McCracken et al., 2023). This lack of mechanistic understanding of Ca^2+^ protective signaling motivated us to study other components of cellular Ca^2+^ handling and signaling, including intracellular stores. However, we found ER-Ca^2+^ levels did not correlate with RGC survival outcome (Supplemental Figure 4), and while mito-Ca^2+^ levels could be correlated with RGC survival, manipulations in accordance with this phenotype were detrimental in both healthy and degenerating RGCs (Figures 4-5). Thus, the precise mechanisms that underlie the protective nature of homeostatic Ca^2+^ signaling in RGC cytosol remain elusive.

It is intriguing that we did not observe a correlation between ER-Ca^2+^ and RGC survival, nor was homeostatic ER-Ca^2+^ altered by ONC. In mouse glaucoma models, Ca^2+^ handling after light stimulation is perturbed in that Ca^2+^ transients have prolonged decay times. This defect is linked to degraded ER-Ca^2+^ buffering and can be corrected with Serca2 overexpression that also protects RGCs from degeneration (Shiga et al., 2024). ER-Ca^2+^ dysregulation is also present in other neurodegenerative disorders including trauma and white matter lesions (Stirling et al., 2014; Villegas et al., 2014), which share some degenerative mechanisms with ONC.

It is perhaps unsurprising that elevating mito-Ca^2+^ promotes RGC degeneration. Mito-Ca^2+^ dysregulation has been implicated in many major neurodegenerative diseases including Alzheimer’s (Ryan et al., 2021; Supnet and Bezprozvanny, 2010), Huntington’s Disease (Jurcau and Jurcau, 2023) and Parkinson’s Disease (PD), which has a strong link between mito-Ca^2+^ dysfunction and neuronal death (Dey et al., 2020; Verma et al., 2017). Mito-Ca^2+^ dysregulation is also observed in axon injury paradigms more closely aligned with RGC degeneration. For example, spinal nerve ligation reduces mito-Ca^2+^ buffering capacity in sensory neurons (Hogan et al., 2014). Injury to spinal cord white matter leads to mitochondrial dysfunction including mito-Ca^2+^ elevation and reactive oxygen species accumulation (Breckwoldt et al., 2014; Villegas et al., 2014). Overall, the more surprising result of our study is the fact that natively well-surviving RGCs are enriched for higher homeostatic mito-Ca^2+^ levels.

Our findings also implicate the MCU complex as a possible target for neuroprotection. MCU has previously been identified as a regulator of multiple neurodegeneration conditions. For example, in a zebrafish model of PD, pharmacological and genetic inactivation of MCU protects dopaminergic neurons and rescues PD behavioral phenotypes in PINK1-KO and pink1 mutant animals (Soman et al., 2017; Soman et al., 2019). Additional studies investigating AD and HD identified mitigations in pathology after MCU knockdown in *Drosophila* and mice (Cai et al., 2022; Twyning et al., 2024). MCU knockdown in mouse traumatic brain injury demonstrates protection from acute degeneration through a mechanism that prevented ROS buildup and reduced iron accumulation (Zhang et al., 2019a). Other mitochondrial Ca^2+^ transporters have also been targeted for neuroprotection. Increases in mito-Ca^2+^ and reduced buffering capacity were observed in a range of zebrafish neurodegenerative disorders, and overexpression of the mitochondrial Na^+^/Ca^2+^ exchanger channel NCLX, which extrudes mito-Ca^2+^, suppressed pathological phenotypes (Twyning et al., 2024). MCU overexpression alone also drives degeneration in other CNS models. Increased mito-Ca^2+^ levels, gliosis and neuronal loss were observed with MCU overexpression in hippocampal neurons *in vivo* and *in vitro* (Granatiero et al., 2019). Our results show that MCU overexpression has detrimental effects on RGC survival after ONC, consistent with observations in other neurodegenerative models (Cai et al., 2022; Granatiero et al., 2019; Twyning et al., 2024). Conversely, shRNA-mediated knockdown of MCU, significantly improved RGC survival. Together, these results are in line with previous observations of neurodegeneration models that evaluated MCU and other mitochondrial ion channels, despite contrasting with the intrinsically high mito-Ca^2+^ levels enriched in well-surviving RGCs.

A possible explanation as to why well-surviving RGCs possess higher homeostatic mito-Ca^2+^ levels, despite this being a degenerative stimulus, is that the persistent stress of high mito-Ca^2+^ requires specialized buffering of Ca^2+^ domains/signaling or that broadly protective pathways are continuously active (Fairless et al., 2019). Many Ca^2+^ sequestering and signaling proteins are differentially expressed across RGCs (Jeon and Jeon, 1998; Krieger et al., 2017; Lee et al., 2010; Yi et al., 2012) and have been implicated in degeneration or survival (Krieger et al., 2017; Kwon et al., 2005; Tran et al., 2019). Hormetic factors linking mitochondrial stress and RGC survival could be related to cell death signaling pathways. For example, expression of the pro-apoptotic gene Bax is not uniform across RGCs in the healthy retina (Isenmann et al., 1997; Napankangas et al., 2003) and appears to be lower in well-surviving ip- and αRGCs, as reported in single cell sequencing datasets (Tran et al., 2019). Importantly, Bax relocation drives RGC death after ONC (Maes et al., 2023) and Bax dosage is positively related to outcome in Bax+/- mutant mice or Bax-/- mutants given AAV-Bax overexpression treatments (Donahue et al., 2020). Protective mitochondrial activity might also be more prevalent in higher mito-Ca^2+^ RGCs. Well-surviving RGCs types display high neural activity (McCracken et al., 2023; Tran et al., 2019), which triggers ROS production resultant from ATP demands and mito-Ca^2+^ transients (Dryanovski et al., 2013; Hongpaisan et al., 2003). As such, a neuron’s antioxidant capacity is tied to its activity (Baxter et al., 2015; Hasel et al., 2015). Multiple treatments to increase the antioxidant capacity of RGCs have been protective in models of glaucoma and ONC (Jiang et al., 2016; Naguib et al., 2023; Wang and Yuan, 2019). Taken together, it is possible that chronically high mito-Ca^2+^ levels lead to multiple protective modifications to prevent oxidative stress or errant apoptotic induction in RGCs.

It is somewhat surprising that we observed slow and prolonged reductions in homeostatic mito-Ca^2+^ that continued over the 14 days after ONC. This contrasts both with observations of mito-Ca^2+^ overload that have been linked to neurodegeneration (Granatiero et al., 2019; Singh et al., 2022; Verma et al., 2017; Zhang et al., 2023) and our previous observations that cytoplasmic Ca^2+^ is relatively stable after ONC (McCracken et al., 2023). It is possible that lowering mito-Ca^2+^ could contribute to RGC degeneration. For example, ATP generation *via* the citric acid cycle and electron transport chain is directly stimulated by Ca^2+^ elevation in mitochondria (Amrapali Vishwanath et al., 2026; Denton, 2009; Fisher et al., 1973; Glancy et al., 2013; Territo et al., 2000), and RGC ATP levels decrease in glaucoma (Quintero et al., 2022). However, we recently reported increasing RGC ATP levels over the days after ONC (Wang et al., 2021). Furthermore, since we observed that mito-Ca^2+^ was similarly decreased in surviving and dying RGCs, it seems mito-Ca^2+^ reductions are unlikely to determine survival outcome. Given that reducing mito-Ca^2+^ by Ru265 treatment or limiting mito-Ca^2+^ permeability by MCU knockdown protected RGCs from ONC, we predict that intrinsically reducing mito-Ca^2+^ could represent a protective response to degenerative signaling.

## Materials and Methods

Experimental model and Subject Details

### Animals

All experimental procedures were performed in accordance with animal protocols approved by the International Animal Care and Use Committee at Washington University in St. Louis Medical School and in compliance with the NIH Guide for the Care and Use of Laboratory Animals. Male and female mice were used equally, dependent on litters and separated in cohorts of 3–5 siblings. For some experiments, control vs. experimental conditions were tested within individual animals. For Cre dependent expression of AAV in RGCs or subsets of RGCs, genetically modified VGlut2-IRES-Cre (028863; Jackson Labs and KCNG4-Cre (029414; Jackson Labs) mice on a C57/B6J background were used). For Trans-Scleral *in vivo* imaging, KCNG4-Cre mice were crossed with B6(Cg)/Rj-Tyrc/c (000058, Jackson Labs) mice for αRGC specific expression in albino mice with non-pigmented sclera. For overexpression experiments, VGlut2-Cre mice were crossed with B6N.Cg-Gt(ROSA)26Sortm1(CAG-EGFP*)Thm/J (032675, Jackson Labs) mice for outer mitochondrial membrane-GFP expression (not analyzed/used). Mice were aged 4 weeks prior to AAV injections and ONC was performed on mice at 6–8 weeks of age. Mice used for mitochondrial-Twitch2b AAV injection experiments that were quantified (Figures 1, 2, 3, SF1, SF2) were all less than 4 weeks of age when injected, and the number of days post viral injection were carefully considered for these experiments. Mice were housed in a barrier facility in Washington University School of Medicine with 12-h light/dark cycles at 21°C.

## Method Details

### Adeno Associated Virus Preparation

To measure cytoplasmic calcium levels, we used a FRET based biosensor, Twitch2b that was cloned into a Cre-dependent AAV expression cassette pAAV-EF1α-FLEX-Twitch2b (McCracken et al., 2023; Thestrup et al., 2014). To localize Twitch2b to mitochondria or endoplasmic reticulum, we cloned in appropriate N-terminal localization sequences from cytochrome c oxidase 8 or the mouse Vh-chain ER-signal peptide respectively, which have previously been used to localize Twitch2b to these organelles in transgenic mice (Witte et al., 2019). pAAV-hSyn-FLEX-MCU-T2A-mCherry was made using standard cloning methods by cloning the MCU coding region (UniProtID: Q3UMR5) into a backbone from pAAV.hSyn.FLEX.FAPdL5-POST-T2A-dTomato.FLEX.WPRE.SV40 (Addgene 105982) where dTomato had been swapped for mCherry. pAAV-U6-shMCU-hSyn-mCherry was created by cloning shRNA targeted to MCU into the AAV-U6-sgRNA-hSyn-mCherry backbone (Addgene 87916). pAAV-hSyn-DIO-mCherry was used as a control (Addgene 50459). All constructs were packaged into AAV2 by The Hope Center Viral Vectors Core at Washington University in St. Louis. Titers of viral preparations ranged from 2 × 10^12^ to 1.0 × 10^13^ GC/mL, as measured by qPCR. Virus was stored in 10 or 20 μL aliquots in a −80°C freezer.

### Intravitreal Injections

Mice were anesthetized by intraperitoneal injection of a ketamine and xylazine cocktail (KX) (10 mg/mL and 1 mg/mL, respectively) in saline at a dose of 10 μL/g body weight. A pulled-glass micropipette was inserted near the peripheral retina behind the ora serrata angled to avoid lens damage. Approximately 1-2 μL of vitreous humor was removed prior to AAV injections. AAV aliquots were centrifuged for at least 30 sec prior to injection to remove air bubbles. Between 1.5 to 2 μL of AAV was injected intravitreally using a Hamilton syringe (80950, Hamilton). Any injections resulting in air within the vitreous or injury to the lens were not included in experiments. An anti-bacterial ophthalmic ointment, Terramycin (Zoetis, NADA #8–763), was applied post-operatively to protect the cornea from dehydration and infection. All animals received subcutaneous Meloxicam (10 mg/mL, 10 μL/g of body weight) as postoperative analgesic. Overexpression constructs were injected in the same manner, and mice were allowed 2-3 weeks for AAV expression and subsequent injury or control analyses. For OX experiments, Cholera Toxin Subunit B (Recombinant) (CTB) Alexa Fluor 488 (ThermoFisher, C22841) was injected at 12 days post ONC injury for assessment of optic nerve regeneration (data not shown).

### Optic Nerve Crush

Optic nerve crush (ONC) was performed as previously described (McCracken et al., 2023). Briefly, mice were anesthetized with KX as described above and the optic nerve was exposed intraorbitally and crushed with fine forceps (Fine Science Tools, Carbon #5, No. 11251-10) for 10 sec at a location approximately 500 μm behind the optic disc. Anti-bacterial eye ointment was applied post-operatively to protect the cornea, and animals received subcutaneous Meloxicam (10 mg/mL, 10 μL/g of body weight) as a postoperative analgesic. Animal health was monitored daily for the first 2 days following ONC and every other day for the duration of the 14-day experiment. For chronic time-lapse experiments, mice received a binocular ONC 4 days after their pre-image was taken. For MCU expression manipulation experiments, binocular ONC was performed 3 weeks after AAV injection. Sham ONC cohorts were included, in which the optic nerve was exposed, forceps were placed around the nerve, but were not closed.

### *In vivo* imaging of mouse retinal ganglion cells

Two-photon trans-pupillary *in vivo* imaging of RGCs was carried out as previously described (McCracken et al., 2023; Wang et al., 2025). In short, a Scientifica Hyperscope was used *for in vivo* image acquisition with a Mai Tai HP 100 fs pulsed laser (Spectrophysics), a Pockels cell, and a pair of galvo mirrors. A 20 mm working distance objective (Mitutoyo, 20X air, 0.4 N.A., 378-824-5) was used with a motorized objective mount for z-stepping. A Chromoflex light collection system paired with GaAsP photo-multiplier tube (PMT) detectors (Scientifica) was used for light collection. A filter cube consisting of a 505 long pass dichroic and 480/40 and 535/30 band-pass filter pairs was used to spectrally filter light for FRET analysis.

Mice were anesthetized with a KX cocktail as described above and placed in an imaging head holder (SGM 4, Narishige). The head of the mouse was secured with pins inserted into the ear canals and a bite bar to hold the maxillary incisors. A solution of 1% w/v atropine and 2.5% w/v phenylephrine hydrochloride in RO water was applied to dilate the pupil, and animals were placed in the dark for 5–15 min prior to imaging. Genteal-tears eye ointment (Alcon Inc.) was applied to both eyes and the head of the mouse was angled to align the iris with the light path. A #1.5 coverslip was placed in a compact filter holder (Thorlabs, DH1) and the holder was fixed to the microscope stage. Using blue LED epifluorescence light to visualize fluorescent cells, the head holder was adjusted to “flatten” the imaging area and reduce signal noise.

Laser power measured out of the objective ranged from 20 to 45 mW and was limited to a maximum of 45 mW to prevent retinal or corneal damage (McCracken et al., 2023; Wang et al., 2025). The Mai Tai pulsed laser was set at 850 nm to most effectively excite the Twitch-2b sensor (Williams et al., 2014). ScanImage acquisition software (Vidreo Technologies) was used to obtain image stacks with an 8 μm z-stepping. Images (512 pixels × 512 pixels, 1 pixel ≅ 1μm^2^) were collected at 0.93 Hz with a 3-frame average. Retinas were scanned from the ganglion cell layer toward the inner nuclear layer to reduce photoreceptor activation. Other imaging settings like PMT voltage, bias voltage, and digital zoom were kept constant across all images.

For chronic-time lapse imaging, VGlut2-Cre mice with Mito-T2b or ER-T2b injections were imaged prior to injury (‘pre-image’). Optimal regions of the retina were identified for chronic imaging, i.e., the region was flat, had robust Twitch2b expression, and could be re-found quickly using vascular landmarks. All pre-images were obtained 100–300 μm away from the optic nerve head at the closest portion of the image. Two or three pre-images were obtained per retina, usually spanning the dorsal-temporal to ventral-temporal retina. In samples with lower Twitch-2b intensity, as occurred in several Mito-T2b or ER-T2b samples, multiple pre-images of the same region were obtained 4-7 days after initial pre imaging. Optic nerve crush injury was performed on both eyes of mice 4 days after the last ‘pre-image’.

Trans-scleral *in vivo* retinal imaging was adapted from previously established trans-scleral imaging approaches (Alarcon-Martinez et al., 2020; Quintero et al., 2022; Takihara et al., 2015). KCNG4-Cre driver lines were bred onto an albino background (B6(Cg)-Tyrc-2J/J; Jackson Lab strain #000058) to remove ocular pigmentation and enable direct trans-scleral imaging. Imaging was performed using a 25× water-immersion objective (NA1.05, working distance 2.0 mm; Olympus). To achieve high spatial resolution of RGC soma and neurites, images were acquired at 2× and 4× digital zoom with a z-step size of 2 µm, corresponding to lateral resolutions of 0.625 µm/pixel and 0.312 µm/pixels respectively. Laser power at the sample was maintained at 10–15 mW to minimize phototoxicity and avoid retinal or RGC damage.

During trans-scleral imaging, mice were maintained under prolonged anesthesia using low-dose isoflurane (0.5% in air, 0.4 L/min), as required. Isoflurane was delivered *via* a custom-designed 3D-printed mouse mask integrated with the head-holding apparatus, enabling stable positioning and bilateral imaging. Supplemental isoflurane was administered approximately 1 hour after imaging onset to maintain an adequate anesthetic plane. Following all imaging sessions, STYE sterile eye lubricant (Prestige Medical) was applied to prevent dry eyes and mice were placed on a 36°C heating pad until they recovered from anesthesia.

### Injections and *in vivo* imaging

For pharmacological injection experiments, drugs were injected intravitreally with a pulled-glass micropipette that was inserted near the peripheral retina behind the ora serrata and deliberately angled to avoid damage to the lens while the mouse was stationed in the headholder. Ru2(μ-N)(NH3)8Cl2]Cl3, trans,trans-Octaamminedichloro-μ-nitridodi-Ruthenium(3+) trichloride, Nitride, ruthenium complex (i.e. Ruthenium Red 265 (Ru265); Sigma, SML2991) was dissolved at a concentration of 2 mM in a 50% Dimethyl Sulfoxide (DMSO; Sigma, 67-68-5) 50% phosphate buffered saline (PBS, made in lab, pH 7.4). Ru265 was aliquoted in DMSO at 10 mM, placed in -20 deg C for long term storage, made fresh weekly, and used for up to 5 five days after preparation while being stored in 4 deg C as a working solution. 200 μM and 20 μM Ru265 solutions were freshly made from 2 mM prepared stocks and used/stored in the same fashion. Prior to injection, drugs were vortexed and sonicated until fully dissolved and centrifuged for ∼10 minutes at 400 rpm to avoid bubbles and precipitate when injecting. Before injection, two pre-images for a single eye were obtained, and animals were prepared for injection on the microscope stage with a modified stereoscope. While still in the imaging head holder, approximately 1.5 μL of solution was injected into the vitreous space, as described above, without vitreous removal. Solutions were injected slowly over 30-60 sec to avoid leakage of fluid from the eye. A post-injection image was obtained 1-2 min after injection, and more post-images were obtained approximately every 3-5 min. Post 2-min samples for 2 mM Ru265 were imaged at 2.1 min ± 8 sec, and post 2-min samples for vehicle control injections (50% PBS, 50% DMSO) were imaged at 2.0 min ± 5 sec. For post 10-min injection analysis, 2 mM Ru265 injected samples were imaged and analyzed at 10.4 ± 1.4 min and vehicle control injections (50% PBS, 50% DMSO) were imaged and analyzed at 9.8 ± 1.22 min. To assess the timing of manipulations, 3 samples for both Ru265 and control conditions were post-imaged for up to two hours; 2 mM concentration of Ru265 had an average time of 83 ± 13.6 min to return to baseline, and vehicle control injections (50% PBS, 50% DMSO) had an average time of 13.5 ± 5.4 min to return to baseline.

### Longitudinal injections and optic nerve crush

Ru265 injections at concentrations of 2 mM, 200 μM, and 20 μM, along with control injections (50% DMSO, 50% PBS), and PBS only were used for longitudinal injection experiments. Another cohort of sham injections was also included, where the eye was punctured but no fluid was injected. Mice were anesthetized and ONC was performed as described above. Injections were performed immediately after ONC (within 2-3 min) as described above, with the only exception that care was made to avoid the part of the eye where the conjunctiva was cut to access the optic nerve. Therefore, injections were performed at the nasal or dorsal portions of the eye rather than temporally. Injections were performed without removing vitreous, and care was taken to avoid the lens. Additional injections were performed every third day after ONC (days 3, 6, 9 and 12 post crush). Eyes that had lens damage or blood in the eye from multiple injections were not included in analysis.

### *In vivo* imaging analysis

Measuring and analyzing *in vivo* Ca^2+^ levels of RGCs was performed as previously described using ImageJ software (McCracken et al., 2023). Raw images were processed and made into two-channel (cpVenus (YFP), pseudo-colored magenta and mCerulean3 (CFP), pseudo-colored green) image stacks. Cell bodies were identified and manually labeled as regions of interest (ROIs) in their brightest and best resolved z-sections. Mean pixel values within each ROI for both CFP and YFP channels were measured with a custom MATLAB script and ratios were recorded. Background was subtracted based on the mode pixel value for each channel.

Variance of the YFP and CFP values within each individual ROI was also calculated. Highly variant RGCs were excluded using a ‘percent error’ value, as determined by integrating pixel intensities for each channel and forming a 95% confidence interval for ratio measurements within RGCs; see McCracken et al., 2023 methods. Any cells with percent error higher than 60% and/or dim RGCs with a raw pixel intensity value (YFP + CFP) lower than 100 were excluded from analysis.

For chronic longitudinal ONC analysis, image stacks were processed by maximally projecting processed images to a single plane, combining the time series of max-projections into a t-stack multi-tiff, and aligning these images with the Linear Stack Alignment with SIFT plugin in ImageJ. ROIs were selected based on cells that could be tracked for all or most of the time-lapse. The mean number of trackable RGCs from a single retina was in mito-T2b experiments was 54 ± 5.8.

For *in vivo* imaging and drug injection experiments, image processing was performed as described above and pre-vs. post-injection images at post-2 min and post-10 min were aligned manually. ROIs were selected for cells that remained in-focus and in-frame for both pre- and post-images. ROIs were circled on the pre-image blind to ratios in the post-images. The change in ratio from pre-image to post-image was quantified as the delta R/R_o_, which was obtained by comparing the ratio of an ROI in the post-image, R_post_, to the ROIs of the same cell in the pre-image, R_pre_ (R_pre_ = R_o_) with the following equation: ΔR/R_o_ = (R_post_ -R_pre_)/R_pre_.

### Histology

All animals were given an intraperitoneal overdose injection of Tribromoethanol (Avertin, 500 mg/kg) (Sigma, T48402) and transcardiacally perfused with ice-cold Phosphate-Buffered Saline (PBS) followed by 100 mL of 4% paraformaldehyde (Sigma) in PBS. Heads were stored in PBS at 4°C, and eyes were removed within 24–48 hr of perfusion, and retinas were dissected and preserved in PBS at 4°C.

After dissection, whole retinas were kept in PBS at 4°C. Retinas were sunk overnight in 30% sucrose in PBS at 4°C and freeze-thawed 3 times using dry ice and a glass slide. Samples were washed with PBS 3 times for 10 min and put in blocking solution (10% normal horse serum (Sigma, 158127) and 0.5% Triton X-100 (Sigma, 11332481001) in PBS) for 1–3 hr at room temperature. Samples were placed in a solution of primary antibodies diluted in blocking solution on a shaker at 4°C for 5–7 days. Primary antibodies used were the following: guinea pig anti-Rbpms (1:2000, Raygene A008712), rabbit anti-MCU (1:500, Cell Signaling, 2250), rabbit anti-TOMM20 (1:500, Proteintech, 11802-1-AP), rat anti-TBR2 (1:500, Invitrogen, 14-4875-82) and goat anti-Osteopontin (SPP1; 1:500; AF808). Retinas were washed with PBS 3 times for 10 min at room temperature. Secondary antibodies (Jackson ImmunoResearch) were diluted in blocking solution for 2–3 d at 4°C. Secondaries were raised in donkey against the primary antibodies host species, cross absorbed and conjugated to Alexa Fluor 405, 488, 568 or 647, and used at 1:500 dilution. After washing three times in PBS, whole retinas were mounted onto glass slides with Vectashield Antifade Mounting Medium (Vector Labs, H-1000-10), cover-slipped, and sealed with nail-polish.

### Confocal microscopy and analysis

For RGC density quantifications (Figures 4-5, SF5, SF7), an inverted laser scanning confocal microscope (Zeiss, Model 710) equipped with a 20× air objective (Zeiss ‘Plan Apochromat’, 0.8 NA, No10098) was used to acquire image stacks of wholemounted retinas at a 1.5 μm z-spacing and 0.446 μm xy pixel resolution. A montage of a 4 X 4 tiled field of the retina surrounding the optic nerve head was obtained, the total region being 2100 μm × 2100 μm, and images were automatically stitched with the Zeiss Blue software. Optic nerves were imaged with the same settings, where the crush site was identified by observing a clear stop in CTB labeling near the end of the optic nerve head. Samples without a clear ONC site were assessed for technical issues, i.e. overcrush or undercrush. Eyes were not included for final analysis if they were determined to be over/under-crushed when comparing the sample’s retina with optic nerve.

For MCU genetic manipulation and longitudinal injection experiments, RGC survival was assessed using RBPMS+ cell density quantification. The cell counter plugin in Fiji was used to count RBPMS+ cells in two representative regions of wholemounted retinas. The size of each region was kept consistent for each experimental cohort, where for Ru2665 injections a size of 101252 μm^2^ was used, for MCU-OX experiments a size of 99670 mm^2^ was used and for shMCU experiments a size of 76593 mm^2^ was used. RGC density values for each individual retina were identified by averaging the RBPMS+ density values in each region.

For *in vivo* imaging and correlative histology experiments, *in vivo* imaged regions were identified on the wholemount retina using vascular and cellular landmarks. Multi-channel images were obtained initially with low resolution to obtain an image overview. Once *in vivo* images were matched to wholemount locations, higher resolution image volumes were acquired using 1 μm z-step, 4x averaging, 0.223 μm xy pixel resolution, and sequential laser scanning.

For assessment of mito-T2b expression in the mitochondria (Figure 1L, Supplemental Figure 3) confocal images were obtained with a 64x oil objective (Zeiss ‘Plan Apochromat’, 1.4 NA, 420782-9900-000). Image stacks were obtained with a 0.25 μm z-step, 0.071 μm xy pixel resolution, and sequential laser excitation were used to avoid crosstalk between signals. Cells were categorized into qualitative groups based on the degree of biosensor mislocalization in the cytoplasm. The vast majority of RGCs displayed clear localization to mitochondria and no mislocalization. RGCs that had mito-T2b mislocalization were either termed complete mislocalization, where cells appeared unhealthy without obvious mitochondrial localization of T2b, or partial mislocalization, where cells had mostly bright mitochondrial localization, but also some signal in the cytoplasm and axons compared to other cells in the imaging field. RGCs were quantified for mito-T2b and TOMM20 intensity values using Z-scores (Mean - (intensity value / standard deviation)), with mean and standard deviation determined within individual high-resolution image regions.

### Quantification and statistical analysis

Statistical analyses were performed using Microsoft Excell and an online publicly available Mann Whitney U test calculator (Social Science Statistics). Details of statistical analyses including number of samples, what sample numbers represent, statistical tests used, and value representations can be found in the Results text and figure legends. Unless otherwise indicated, values are mean ± S.E.M.

### Data availability

All data are available form the corresponding author upon request.

### Additional information

We thank Michael Casey and Mingjie Li for assistance. This work was supported by the Hope Center Viral Vectors Core at Washington University School of Medicine, an unrestricted grant (to the Department of Ophthalmology and Visual Sciences) from Research to Prevent Blindness, and Vision Core Grant P30 EY002687.

### Author contributions

S.M., M.Z. and P.R.W. designed experiments. S.M., K.S., M.Z., C.Z., S.B.T and M.A. conducted experiments. S.M. and P.R.W. wrote the original draft. All authors contributed to revising the final version of the manuscript.

### Funding

National Eye Institute (NEI) Institutional National Research Service Award T32 EY013360

-Sean McCracken

Research to Prevent Blindness Career Development Award

-Philip R. Williams

BrightFocus Foundation National Glaucoma Research

-Philip R. Williams

Alcon Research Institute Young Investigator Award

-Philip R. Williams

NEI grants EY032908, EY035684, EY036111, EY012543

-Philip R. Williams

This work was supported by an unrestricted grant to the Department of Ophthalmology and Visual Sciences from Resarch to Prevent Blindness, and by Vision Core Grant P30 EY002687

## Supplemental Figure Captions

**Supplemental Figure 1. Ru265 decreases Mito-Ca^2+^ levels in a dose dependent manner.** (A) Example *in vivo* 2-photon aips of mito-T2b before and 10 min after injection of vehicle control (50% DMSO and 50% PBS). (B) Line graphs of mito-Ca^2+^ levels before and 10 min after vehicle injection. Individual RGCs are in grey, and mean is in black, MWU test (n = 95 RGCs from 3 retinas and 3 mice). (C) Swarm plot of changes in mito-Ca^2+^ levels after vehicle injection. (D) Example *in vivo* 2-photon aips of mito-T2b before and 10 min after injection of 20 μM Ru265. Arrow indicates RGC with reduction in mito-Ca^2+^. (E) Line graphs of mito-Ca^2+^ levels before and 10 min after 20 μM Ru265 injection. Individual RGCs are in grey, and mean is in black, MWU test (n = 101 RGCs from 4 retinas and 4 mice). (F) Swarm plot of changes in mito-Ca^2+^ levels after 20 μM Ru265injection. (G) Example *in vivo* 2-photon aips of mito-T2b before and 10 min after injection of 200 μM Ru265. Arrows indicate RGCs with reduction in mito-Ca^2+^. (H) Line graphs of mito-Ca^2+^ levels before and 10 min after 200 μM Ru265 injection. Individual RGCs are in grey, and mean is in black, MWU test (n = 159 RGCs from 4 retinas and 4 mice). (I) Swarm plot of changes in mito-Ca^2+^ levels after 200 μM Ru265 injection. Scale bars = 50 μm.

**Supplemental Figure 2. Ru265 induces Ca^2+^ elevations at 2 minutes post injection, likely due to DMSO vehicle.** (A) Example *in vivo* 2-photon aips of mito-T2b before and 2 min after injection of 2 mM Ru265 (left). Line graphs of mito-Ca^2+^ levels before and 2 min after 2 mM Ru265 injection. Individual RGCs are in blue, and mean is in black (n = 154 RGCs from 4 retinas and 4 mice). (B) Example *in vivo* 2-photon aips of cyto-T2b before and 2 min after injection of 2 mM Ru265 (left). Line graphs of cyto-Ca^2+^ levels before and 2 min after 2 mM Ru265 injection. Individual RGCs are in orange, and mean is in black (n = 96 RGCs from 4 retinas and 4 mice). (C) Swarm plots comparing changes in Ca^2+^ levels 2 min after 2 mM Ru265 between cyto- and mito-T2b. Black bars are mean +/- SEM, MWU test. (D) Example *in vivo* 2-photon aips of mito-T2b before and 2 min after injection of DMSO/PBS vehicle (left). Line graphs of mito-Ca^2+^ levels before and 2 min after vehicle injection. Individual RGCs are in blue, and mean is in black (n = 73 RGCs from 3 retinas and 3 mice). (E) Example *in vivo* 2-photon aips of cyto-T2b before and 2 min after injection of behicle (left). Line graphs of cyto-Ca^2+^ levels before and 2 min after vehicle injection. Individual RGCs are in orange, and mean is in black (n = 87 RGCs from 3 retinas and 3 mice). (F) Swarm plots comparing changes in Ca^2+^ levels 2 min after vehicle injection between cyto- and mito-T2b, MWU test. Scale bars = 50 μm.

**Supplemental Figure 3. Mitochondrial targeted Twitch2b mislocalization occurs after longer expression durations.** (A) Example confocal mips of fixed retinal wholemount showing endogenous mito-T2b expression (green) and TOMM20 immunostaining (magenta) at the indicated time points after AAV injection. Yellow arrow indicates RGC with partial mislocaliztion and orange arrows indicate RGCs with complete mislocaliztion of mito-T2b. (B) Example confocal mips of ‘healthy’ RGCs expressing mito-T2b co-localized with TOMM20, co-stained with RBPMS (blue). (C) Example confocal mips of RGCs expressing mito-T2b that is ‘partially mislocalizated’ and immunostained for TOMM20 and RBPMS. (D) Example confocal mips of RGCs expressing mito-T2b that is ‘completely mislocalizated’ and immunostained for TOMM20 and RBPMS. (E-F) Pie charts showing the proportion of RGCs in each of the qualitative mito-T2b localization groups at indicated time points after AAV injection (< 30 dpi n = 917 RGCs from 5 retinas and 3 mice; > 60 dpi n = 446 RGCs from 4 retinas from 4 mice mice). (G) Percentage of RGCs in indicated mislocalization groups at indicated time points. Graph is sample means +/-SEM (MWU test). (H-I) Z-score of mito-T2b expression intensity measure from fixed wholemounts representing indicated qualitative localization groups and time points after AAV injection. Graph is sample means +/- SEM. Scale bars = 10 μm.

**Supplemental Figure 4. Endoplasmic Reticulum Ca^2+^ levels are not correlated with survival to ONC.** (A) Example *in vivo* 2-photon mips of cytoplasmic cyto-T2b (left) and ER-2b (right) expressed in RGCs of VGlut2-Cre mice. (B) Swarm plots of homeostatic cyto-, mito-, and ER-Ca^2+^ levels, as measured by cpVenus (YFP) to mCerulean (CFP) FRET ratios (cyto-T2b and mito-T2b replotted from Figure 1; ER-T2b = 717 RGCs from 9 retinas and 5 mice. MWU test). (C) Example *in vivo* 2-photon aips of ER-Ca^2+^ expressing RGCs before ONC and at the indicated days post ONC (dpc). (D) Swarm plots of pre-injury ER-Ca^2+^ levels in RGCs that survived (green) or died (red) following ONC. Means +/- SEM are labeled with black bars (survive n = 28 RGCs from 4 retinas and 3 mice, die n = 133 RGCs. MWU test). Scale bars = 100 μm.

**Supplemental Figure 5. Chronic Ru265 treatment does not kill RGCs.** (A) Representative mips of retinal wholemounts immunostained with RBPMS at 14 days after sham ONC and indicated treatments. (B) Quantification of RBPMS densities from indicated experimental conditions. Points are individual retinas bars are mean +/- SEM (Sham ONC + Vehicle n = 5 retinas from 4 mice; Sham ONC + Sham injection n = 4 retinas from 4 mice; Sham ONC + Ru265 n = 5 retinas from 4 mice). (C) Representative mips of retinal wholemounts immunostained with RBPMS at 14 days after ONC and indicated sham treatments. (D) Quantification of RBPMS densities from indicated experimental conditions. Points are individual retinas bars are mean +/- SEM (ONC + Sham n = 6 retinas from 6 mice; ONC + PBS n = 5 retinas from 5 mice; ONC + Vehicle n = 6 retinas from 6 mice). Scale bars = 100 μm.

**Supplemental Figure 6. Validation of AAV mediated manipulation of MCU expression.** (A) Example confocal mips of retinal wholemounts immunostained for MCU (green) and RBPMS (blue) with endogenous mCherry reporter signal from AAV-MCU-OX treatment (red). (B) Scatterplot of MCU immunofluorescence intensity and mCherry reporter signal measured from confocal wholemount images. Dashed line represents trendline for all RGCs with a 95% confidence interval. Grey background shading indicates threshold for flagging RGCs as mCherry- or mCherry+. (C) Box and whisker plaots of MCU immunostaining intensity compared between MCU-OX mCherry reporter positive and negative RGCs. Points are individual RGCs, boxes depict the lower quartiles, the median, and the upper quartiles while the whiskers represent the minimum and maximum datapoint at 1.5 x the interquartile range. MWU test (n = 504 RGCs from 3 retinas and 3 mice). (D) Example confocal mips of retinal wholemounts immunostained for MCU (green) and RBPMS (blue) with endogenous mCherry reporter signal from AAV-shMCU treatment (red). (E) Scatterplot of MCU immunofluorescence intensity and mCherry reporter signal measured from confocal wholemount images. Dashed line represents trendline for all RGCs with a 95% confidence interval. Grey shading indicates threshold for flagging cells as mCherry- or mCherry+. (F) Box and whisker plots of MCU immunostaining intensity compared between shMCU mCherry reporter positive and negative RGCs. Points are individual cells, boxes depict the lower quartiles, the median, and the upper quartiles while whiskers represent the minimum and maximum datapoint at 1.5 x the interquartile range, MWU test (n = 489 RGCs from 3 retinas and 3 mice). Scale bars = 100 μm.

**Supplemental Figure 7. MCU-OX leads to slow RGC degeneration.** (A) Representative mips of fixed retinal wholemounts immunostained for RBPMS at 14 days after sham surgery. (B) Quantification of RBPMS+ cell density in mice with intravitreal injections of indicated AAV MCU gene expression manipulations compared to littermate AAV-mCherry treated controls. Points indicate individual eyes, bars indicate mean +/- SEM. (mCherry n = 6 retinas from 6 mice, shMCU n = 3 retinas from 3 mice, MCU-OX n = 7 retinas from 7 mice). (C) Representative mips of fixed retinal wholemounts immunostained for RBPMS at 8 weeks after sham surgery. (D) Quantification of RBPMS+ cell density in mice with intravitreal injections of indicated AAV MCU gene expression manipulations compared to littermate AAV-mCherry treated controls. Points indicate individual eyes, bars indicate mean +/- SEM, MWU test. (mCherry n = 4 retinas from 4 mice, shMCU n = 3 retinas from 3 mice, MCU-OX n = 3 retinas from 3 mice). Scale bars = 100 μm.

